# Century-old chromatin architecture preserved with formaldehyde

**DOI:** 10.1101/2023.07.26.550239

**Authors:** Erin E. Hahn, Jiri Stiller, Marina Alexander, Alicia Grealy, Jennifer M. Taylor, Nicola Jackson, Celine H. Frere, Clare E. Holleley

## Abstract

Co-ordinated regulation or dysregulation of chromatin architecture underpins fundamental biological processes, such as embryonic development, disease, cellular programming and response to environmental stress. The dynamic and plastic nature of chromatin accessibility is a major driver of phenotypic diversity, but we know shockingly little about the temporal dynamics of chromatin reorganisation and almost nothing prior to the existence of flash-frozen specimens. Linking two disparate fields by their common use and application of the preservative formaldehyde, we present an approach to characterise chromatin architecture in formaldehyde-preserved specimens up to 117 years old. We characterise how over-fixation modulates but does not eliminate genome-wide patterns of differential chromatin accessibility. Our novel analytical approach identifies promoter regions enriched for gene ontology terms matching the tissue of origin, resulting in sex-specific and environment-dependent genome-wide profiles. Contrary to prevailing dogma, we show that over-fixation is essential for the successful recovery of historical chromatin architecture. Our methodological and analytical advances open the door to the first detailed and comprehensive view of the epigenetic past and reveal a new role for museum collections in understanding chromatin architecture dynamics over the last century.

## Main

Chromatin, the cell’s intricate web of DNA and proteins, orchestrates gene expression, shaping cellular identity and dynamically responding to ever-changing signals. Chromatin compaction exists on a spectrum from tightly packed (typically transcriptionally silent) heterochromatic regions to the more open and accessible (typically transcriptionally active) euchromatic regions. Chromatin architecture therefore provides clues about which genes are being expressed at a given time or set of environmental conditions [1,2]. Characterising chromatin architecture changes throughout life and tracking predictable plastic responses to environmental stressors is an approach poised to reveal functional regulatory mechanisms involved in development, aging, disease and perhaps even the regulatory repertoire of responses to changing climates. However, deep temporal insight into chromatin architecture (decades-centuries) is limited by a mismatch between current molecular capability (e.g.[3]) and the state of preservation in historical specimens [4]. Here we solve this problem by combining the strengths of modern chromatin biology with museum genomics. In this study, we have utilised the common application of formaldehyde fixation in both fields to generate new epigenomic capability and deep temporal insight into chromatin architecture.

The field of chromatin biology was born in the mid-20th century, with methods relying upon formaldehyde to preferentially cross-link histone-associated DNA and isolate chromatin. Formaldehyde is still essential in modern techniques such as Micrococcal Nuclease (MNase) [5,6] treatment, Formaldehyde Assisted Isolation of Regulatory Elements (FAIRE) [7], Chromatin Immunoprecipitation (ChIP) [8], and High-throughput Chromosome Conformation Capture (Hi-C) [9]. Similarly, from the early 1900’s, the use of a formaldehyde-based media called formalin (3.7% formaldehyde), came into common use in histopathology, anatomy, and embalming human remains. Formalin media was also used extensively by early naturalists to preserve voucher specimens, facilitate detailed anatomical descriptions and to document the biodiversity of local and explored regions. Thus, many of the earliest collected vertebrate specimens (including taxonomic “type” specimens) have been exposed to formaldehyde. Formaldehyde preservation is most common amongst taxa that do not have alternative means of preservation (e.g., fish, amphibians, reptiles), but is also applied regularly to all biota. Museum preservation practices present several obstacles for molecular applications: 1. Variable rates of specimen decomposition; 2. Uneven fixation during formaldehyde penetration; 3. Significantly heavier fixation of tissue (3.7% versus 1%) for longer periods of time (days versus minutes) and specimens may remain exposed to formaldehyde indefinitely. Thus, a custom approach was required to contend with the unique combination of extreme fixation and significant DNA degradation in old specimens.

To assess gene-regulation spanning the last century, we adapted two chromatin accessibility assays, FAIRE-Seq and MNase-Seq, for use in museum specimens. Specifically, FAIRE-Seq enriches for open chromatin by using phenol chloroform to separate formaldehyde crosslinked protein-associated DNA (e.g., heterochromatin) from unbound regions (e.g., euchromatin). Whereas MNase-Seq enriches for nucleosome-bound DNA through enzymatic digestion of euchromatin. We hypothesised that chromatin architecture is preserved in formaldehyde-exposed historical specimens, observable as sequence read depth variation associated with chromatin accessibility. Preliminary reports hinted at this potential, due to read-periodicity patterns observed in shotgun whole genome sequencing data from formaldehyde-preserved museum specimens [10] that resembled a similar signature observed in ancient DNA [11]. When optimised, archival FAIRE and MNase assays could offer species-agnostic antibody-free assay of historical chromatin accessibility across eukaryotes.

We conducted initial optimisation using a fixation time-series in a well-characterised experimental yeast system (*Saccharomyces cerevisiae*). This series established the molecular consequences of over-fixation on visualising chromatin accessibility. We cultured yeast under optimal and heat shock conditions, measured expression differences from fresh cells via RNA-Seq and processed cells fixed with 1% formaldehyde for 15 minutes, 1 hour, 6 hours and 24 hours with established FAIRE [12] and MNase [13] workflows. Then, we called accessibility signals as occupancy values in DANPOS3 [14] and tested for significant peak width changes (FDR < 0.05) between assay and input control. We observed an assay-specific progressive shift in the abundance and morphology of occupancy signal using both FAIRE and MNase methods (Figures 1A & S1A). Fixation-induced changes in occupancy morphology was most evident in the MNase assay, which also had a higher reproducibility across three technical replicates (genes shared between replicates: MNase = 77 – 83%; FAIRE = 7 – 21%; Figures 1B & S1B). This difference in assay sensitivity and reproducibility is consistent with modern studies that show that FAIRE consistently has a low signal-to-noise ratio compared to other assays [12,15]. Interestingly, fixation time had no significant impact on the proportion of genes shared between replicates in either assay, indicating that fixation alters but does not destroy occupancy signals (Figure S1C).

**Figure 1.**
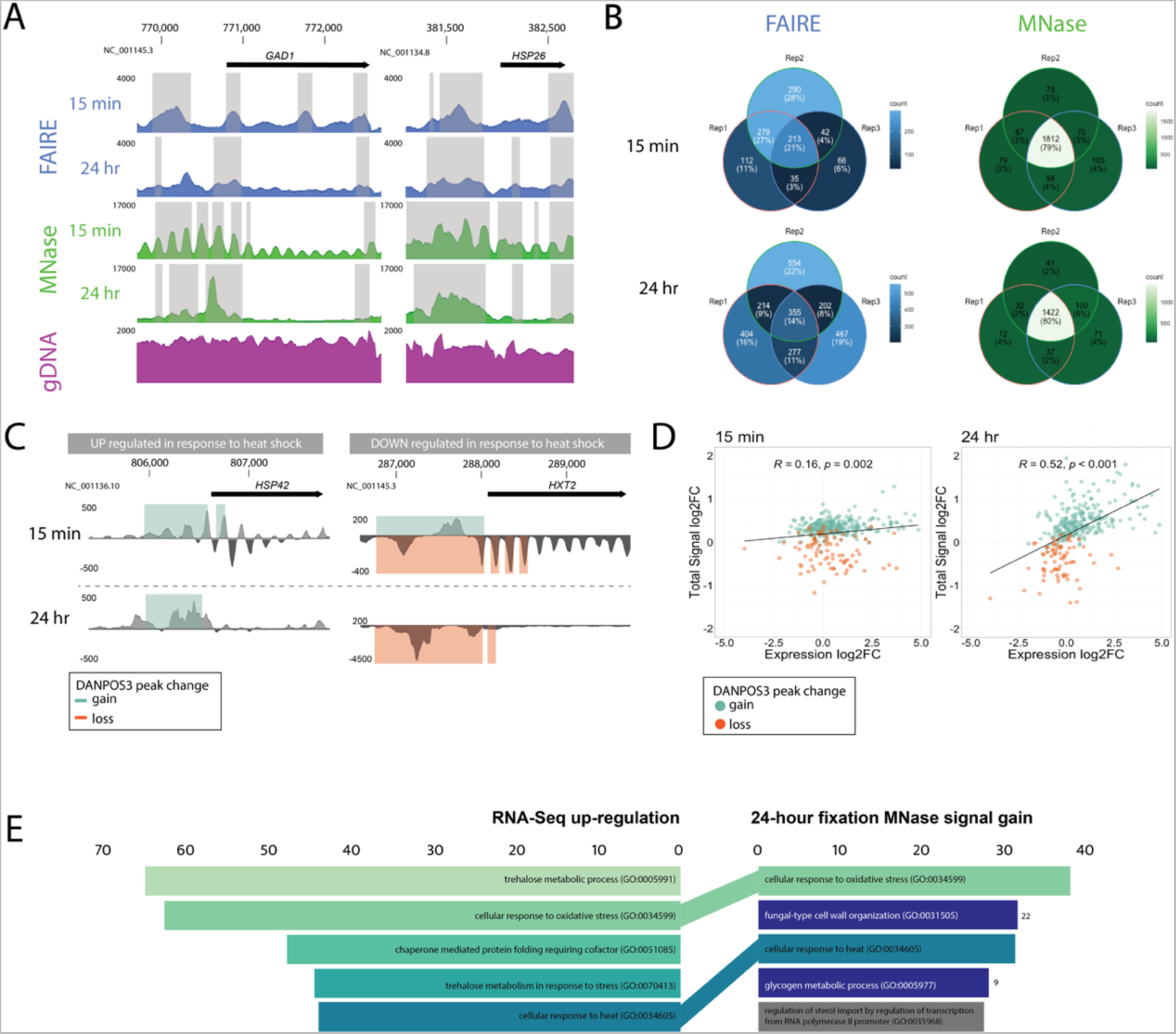
Heavy formaldehyde fixation modulates but does not eliminate chromatin architecture evidence in experimental yeast cultures. **A.** Pooled occupancy values (FAIRE: blue, MNase:green) compared to gDNA extraction control (purple) with formaldehyde fixation for 15 min or 24 hr of heat shocked *S. cerevisiae*. Shading indicates regions with significant peak width shifts (FDR < 0.05) between treatment and input control. Upstream of highly upregulated GAD1 and HSP26 genes (log2FC = 3.8 and 9.02), changes in occupancy signal morphology are observed. The 5’ FAIRE peak broadens, while the distinct 5’ MNase nucleosome array transforms into a single peak. **B.** Venn diagrams demonstrate repeatability of the FAIRE (blue) and MNase (green) assays among technical replicates in heat-shocked yeast cultures fixed with formaldehyde. Numbers/proportions represent genes with significant peak gain (FDR < 0.05, log10Pval < −6) within 2 kb upstream of the TSS. Lighter colors indicate higher shared gene count. **C.** Differential DANPOS3 occupancy values comparing pooled replicates of heat-shocked yeast to optimal growth conditions treated with MNase. Signal changes are shown for a highly up-regulated gene (HSP42, log2FC = 4.86) and a highly down-regulated gene (HXT2, log2FC = −2.21) as measured by RNA-Seq analysis of fresh cultures. Green and orange shading indicate regions of significant (FDR < 0.05, log10Pval < −6) peak gains or losses. **D.** Total signal log2FC for genes with significant (FDR < 0.05) total peak signal change between pooled replicate heat shock and optimal growth conditions in the 2 kb region upstream of the TSS is plotted against expression log2FC measured by RNA-Seq. Only genes detected with significant total peak change at both the 15-min and 24-hour time points are shown. Genes are coloured green for signal gain or orange for signal or loss. Linear regression lines are fitted, and correlation coefficients (R) and p-value are provided for each time point. **E.** GO Biological Process enrichment: Genes with significant peak gain across pooled replicates (FDR < 0.05, log10Pval < −6) within 2 kb upstream of the TSS in MNase-treated yeast fixed for 24-hours (N=383) compared to significantly up-regulated genes (N=352) measured by RNA-Seq. Length of colored bars corresponds to the Enrichr combined score [log(p-value) * z-score]. MNase GO terms are colored a shade of green, dark blue or grey if they are found in the top 5, top 25 or not within the RNA-Seq GO terms.

Genome-wide chromatin accessibility profiles were informative about the regulatory response of yeast to heat-shock under all fixation conditions but was most definitive at the two extremes (15 min versus 24 hours fixation). Using MNase, chromatin occupancy shifts successfully identified the directionality of expression changes in response to heat shock (established by RNA-seq) (Figure 1C) and GO term enrichment analysis with EnrichR [16,17] identified terms associated with heat stress (Figure 1E). Two of the top five GO terms identified in heavily fixed chromatin were shared with terms identified via RNA-Seq (Figure 1E). Additionally, under maximal fixation conditions (24 hours), the MNase assay displayed a significant positive correlation between the magnitude of occupancy shifts and the magnitude of transcriptional activity (R = 0.52, p < 0.001; Figures 1D & S2). Thus, over-fixation of chromatin may provide semi-quantitative information about gene expression, an advance over-and-above the existing utility of modern non-quantitative assays.

Having established that the chromatin accessibility state is recoverable, despite excessively long fixation conditions in yeast, we then adapted both assays for use in heavily fixed archival vertebrate tissues based on established protocols for fresh vertebrate tissues [12,18]. Significant optimisation was required to dissociate fixed multicellular tissue, improve the recovery of highly degraded and heavily crosslinked chromatin, and thus enhance the modulated archival chromatin architecture signal (Figure 2). To robustly develop and test our archival assays under standardised conditions, we created an experimental collection of formaldehyde-preserved inbred C57 Black 6 laboratory and outbred wild-trapped mice (*Mus musculus*) with specimen-matched flash-frozen liver tissue (Table 1).

**Figure 2.**
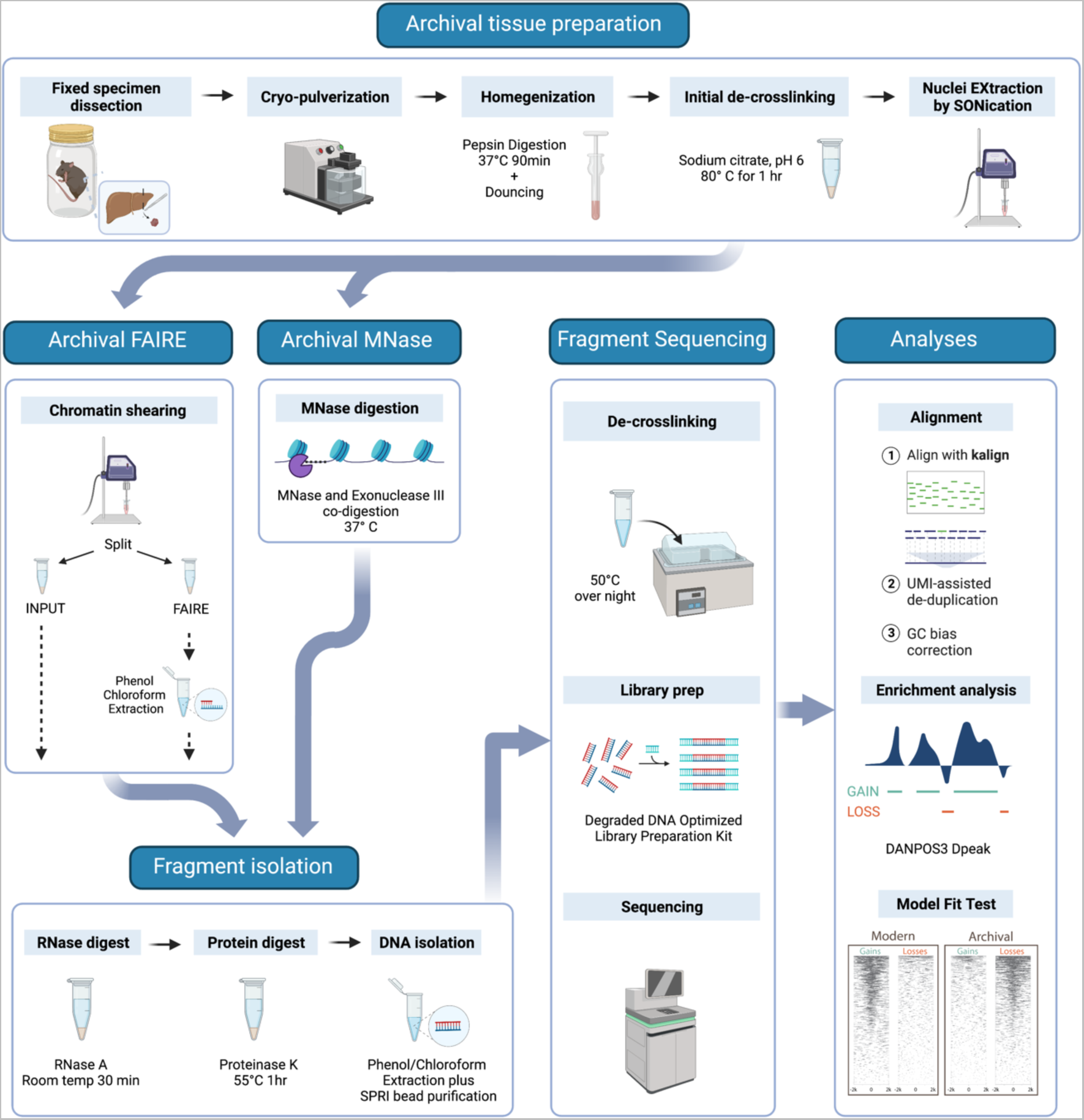
Overview of the archival chromatin assay workflow. We prepare heavily fixed archival tissue nuclei for chromatin extraction through a step-wise process. This includes cryo-pulverization for tissue fracturing, enzymatic digestion with pepsin to improve dissociation, dounce homogenization for fine tissue disruption, and prolonged sonication (NEXSON) for nuclei extraction (modified from Arrigoni et al., 2016). The tissue can then be processed via: FAIRE treatment with further sonication to shear the chromatin followed by reservation of a fraction for input control and phenol:chloroform extraction of the FAIRE fraction or MNase treatment of the nuclei with co-digestion with MNase and Exonuclease III to generate fragments of approximately 100bp. Isolated chromatin then undergoes RNase and proteinase K treatment before DNA fragments are purified using phenol:chloroform extraction and SPRI bead purification optimized for small fragment recovery. Sequencing libraries are prepared using an IDT xGEN cfDNA & FFPE DNA kit for paired-end sequencing. Sequencing reads are mapped using kalign without prior trimming. Alignments are de-duplicated using unique molecular identifiers (UMIs) and undergo GC-bias correction. Enrichment analyses are performed using the DANPOS3 dpeak function. Prior to downstream analysis, confirmation of the expected peak loss versus peak gain pattern can be performed. Refer to the Supplement for detailed methods.

**Table 1.** Specimen details for mock-preserved mouse specimens. We processed six *Mus musculus* individuals for inclusion in a mock-preserved experimental specimen set. For each specimen, we provide the Australian National Wildlife Collection (ANWC) Registration number, the strain or collection site as the Genetic Background, the Time Interval in years between preparation & fixed-tissue sampling, and the individual’s sex, weight, and length.

| Sample name | ANWC Reg. No. | Genetic Background | Time Interval<br>(years) | Sex | Weight<br>(g) | Length<br>(cm) |
| --- | --- | --- | --- | --- | --- | --- |
| Lab1 | M37810 | C57BL6 | 4.8 | M | 23.5 | 9.4 |
| Lab2 | M37811 | C57BL6 | 4.8 | M | 22.7 | 9.5 |
| Lab3 | M37816 | C57BL6 | 4.8 | M | 23.6 | 9.7 |
| Wild1 | M37959 | Wild - Murrumbateman | 2.3 | M | 14.0 | 7.5 |
| Wild2 | M37971 | Wild - Murrumbateman | 2.3 | F | 19.5 | 8.5 |
| Wild3 | M37976 | Wild - Yass | 2.3 | F | 20.0 | 8.5 |

By exploiting the properties of over-fixation, our novel MNase and FAIRE protocols successfully enriched for occupancy signal changes in regions ±2 kb of TSSs (Figures 3A, 3C & S3A-B) that produced occupancy profiles matching to the tissue of origin (Figures 3E & S3C). Notably, the MNase assay of archival laboratory mice tissue signal showed stronger enrichment globally at TSSs compared to FAIRE (Figure S3A), a higher degree of repeatability between replicates (53% overlap in genes identified in all three replicates compared to 31%; Figure S3B), and the MNase gene sets more closely resembled those from fresh tissue compared to the FAIRE assay (51% agreement compared to 38%; Figure S3C). Here, we focus in greater depth upon the archival MNase assay results, however, further exploration of the relative sensitivities of the MNase and FAIRE assays as applied to archival tissues may reveal additional insight into the effects of archival fixation on chromatin architecture as well as providing parallel lines of evidence to characterise historical gene regulation.

**Figure 3.**
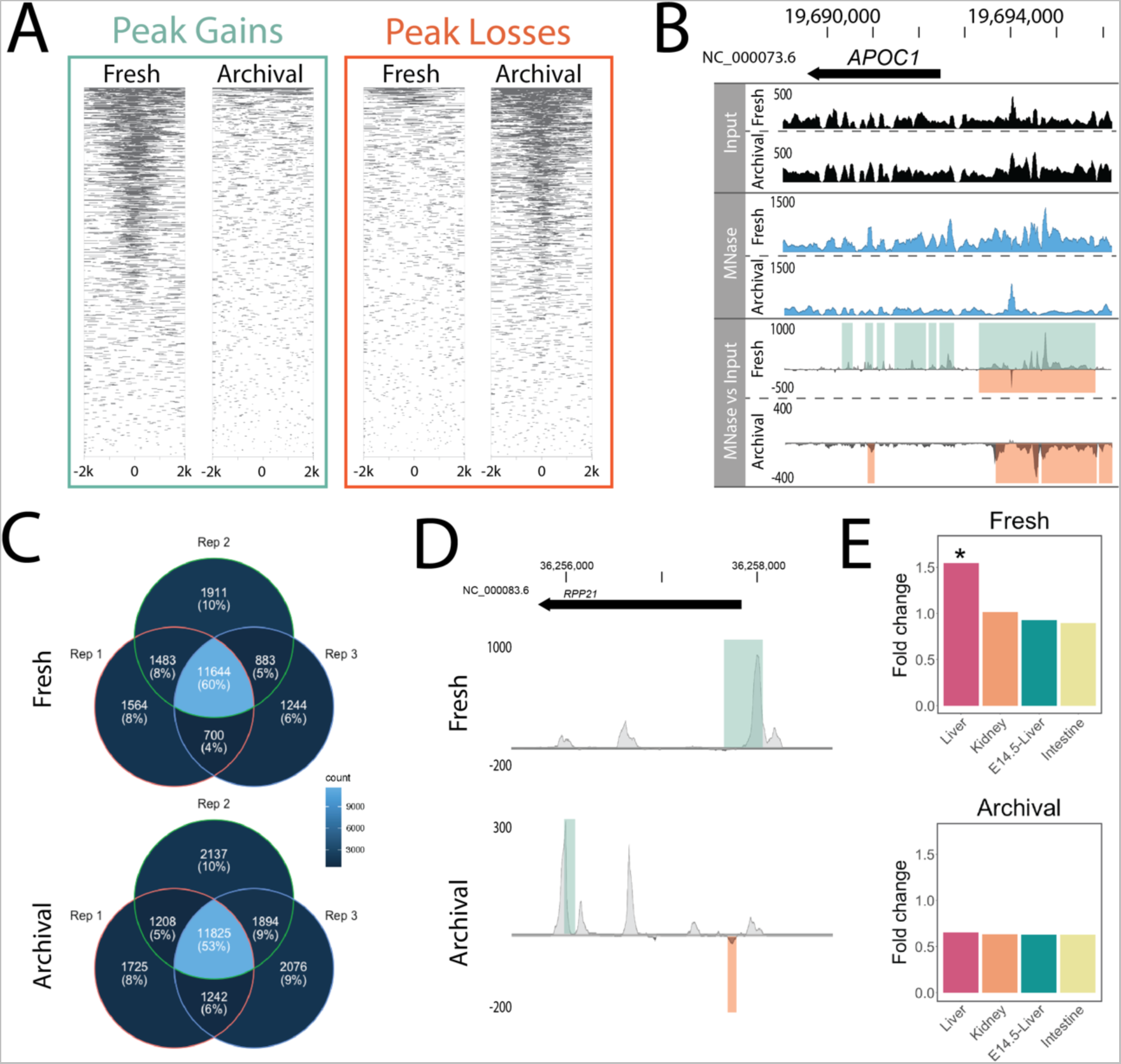
Genome-wide occupancy profiles using MNase in archival mouse specimens are the inverse of freshly collected specimens. **A.** Heatmap of MNase assay significant peak gains and losses (FDR < 0.05, log10Pval < −6) in fresh and archival tissues 2 kb either side of genome-wide transcription start sites pooled across three individuals. **B**. Pooled occupancy values as wiggle traces (DANPOS3 dpeak function) for input (black) and MNase (blue) as well as differential MNase signal over input control (grey) for fresh and archival *Mus musculus* liver tissue. Occupancy values and signal changes are shown upstream of a gene highly expressed in liver (APOC1, FPKM = 38,660) as measured by RNA-Seq analysis of fresh tissue. Green and orange shading upon the differential signal panel represent significant (FDR < 0.05, log10Pval < −6) peak gains or losses across three individuals detected by DANPOS3. **C.** Venn diagrams demonstrate relative repeatability of the MNase assay applied to fresh and archival liver tissue among biological replicates in laboratory mice. Numbers/proportions represent genes with significant peak gains for fresh tissue and losses for archival tissue (FDR < 0.05, log10Pval < −6) within 2 kb upstream of the TSS. Lighter colors indicate higher shared gene count. **D.** Differential pooled DANPOS3 occupancy values comparing laboratory strain to wild caught mice in fresh and archival liver tissue treated with MNase. Signal change is shown for a gene highly upregulated in laboratory versus wild mice (RPP21, log2FC = 9.611) as measured by RNA-Seq analysis of fresh tissue. Green and orange bars represent significant (FDR < 0.05, log10Pval < −6) peak gains or losses detected by DANPOS3. **E.** Genes with pooled occupancy signal changes (Fresh = gains; Archival = losses) show highest enrichment (fold change over a set of all mouse protein coding genes) for genes expressed in liver in both fresh and archival mouse tissues. For each panel, shared gene lists were assembled from a pool of three laboratory and three wild mice and enrichment within Mouse ENCODE datasets was calculated with TissueEnrich [45]. * significant (Benjamin & Hochberg adjusted p-value < 0.001) enrichment above background.

Critically, the genome-wide signature of chromatin architecture in aged multicellular museum specimens (as opposed to yeast cultures), is the inverse of the standard MNase and FAIRE profiles from minimally fixed fresh-tissue (Figures 3A & S3A). This indicates that the archival assays work to reveal historical chromatin accessibility through depletion of open active chromatin rather than through enrichment, a unique property of the archival assay. To explain the inverse occupancy signal, we focus on MNase treatment and propose a model under which fixation, cellular dissociation, and age of specimen influence chromatin accessibility and thus regional enrichment or depletion (Figure 4). This model unifies our mouse and yeast observations and demonstrates that the archival MNase assay is informative across single and multicellular eukaryotes.

**Figure 4.**
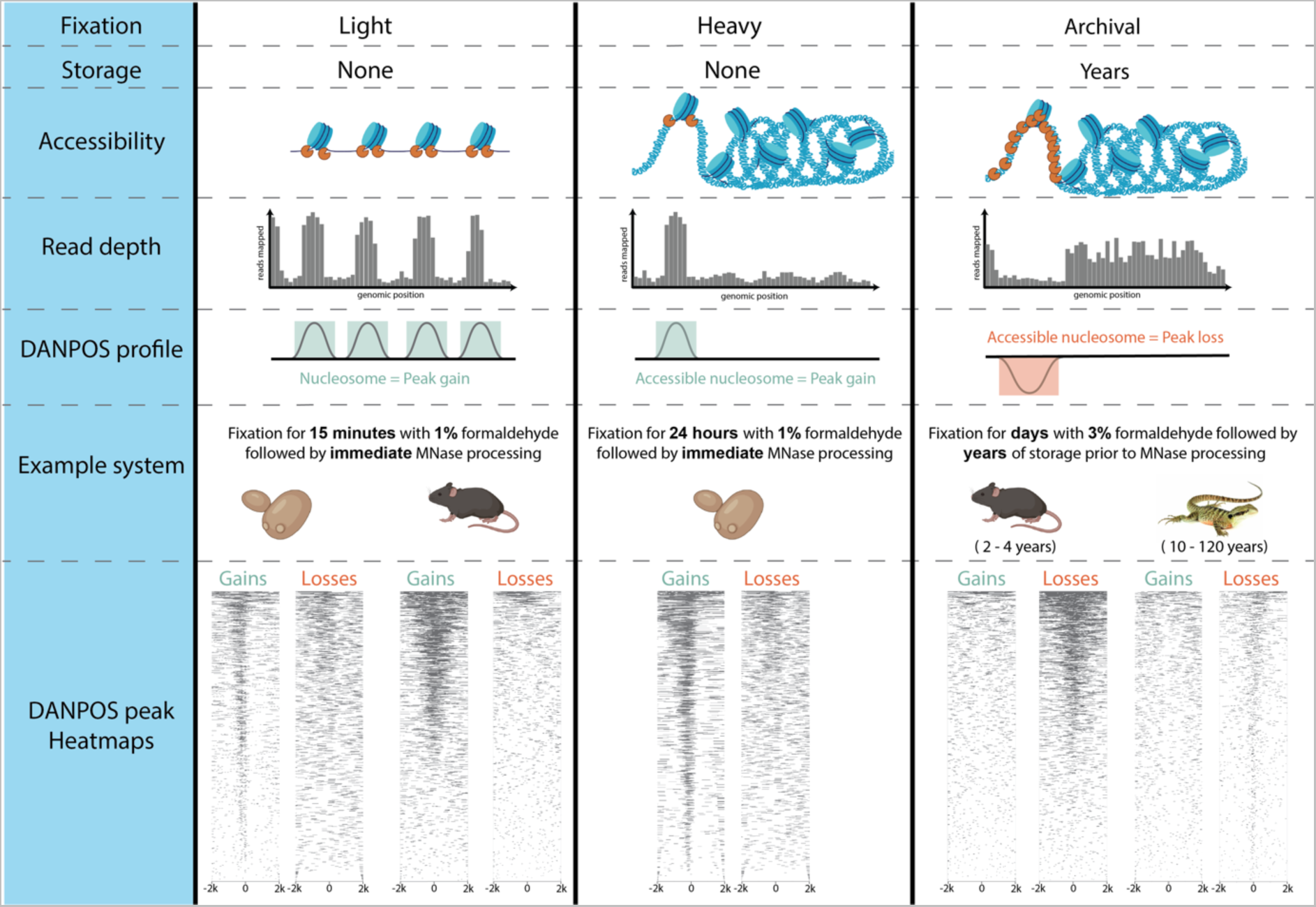
Proposed model of the effect of fixation and long-term storage on MNase accessibility and occupancy signal. Conceptual model of the combined effects of fixation and storage conditions on MNase occupancy signal. From top to bottom, **(Accessibility)** Applied to lightly fixed chromatin, the MNase enzyme (depicted in orange) cleaves DNA adjacent to the nucleosome and resects unbound DNA, thus releasing nucleosome bound DNA. Applied to heavily fixed chromatin, the MNase enzyme’s access to unbound DNA is modulated by chromatin accessibility, thus reducing release of nucleosome-bound DNA only from within the most assessable chromatin regions. Applied to archivally fixed chromatin, prolonged MNase digestion is required to release sufficient DNA for sequencing from the heavily fixed chromatin within intact whole specimens stored for months to many years. This prolonged digestion preferentially degrades both linker DNA and nucleosome-bound DNA in MNase accessible regions and releases fragments from relatively inaccessible regions. **(Read depth)** Relative accessibility of the MNase enzyme alters the read depth pattern observed in region of euchromatin relative to heterochromatin. **(DANPOS profile)** DANPOS efficiently detects both relative occupancy value gains and losses resulting from MNase digestion. **(Example system)** We offer examples of light fixation in both a single (yeast) and multicellular (mouse) system with no storage time, heavy fixation in single (yeast) cellular system with no storage time and archival fixation in two multicellular vertebrate systems stored for several years (mouse) or up to 120 years (water dragon). **(DANPOS peak Heatmaps)** For each example system, we show a heatmap of MNase assay significant peak gains and losses (FDR < 0.05, log10Pval < −6) 2 kb either side of genome-wide transcription start sites pooled across all replicates (three for yeast and mouse, 5 for water dragon). Under light and heavy fixation, the predominant genome-wide signal appears as occupancy gains while under archival fixation, the predominant genome-wide signal appears as occupancy losses irrespective of storage time.

Similar to the 24 hour-fixed yeast data, the magnitude of occupancy change in our archival MNase assay appears to be a semi-quantitative proxy for gene expression (Figures 3B & S4). Highly expressed liver-specific genes (e.g., APOC1, Figure 3B) had a strong depletion signal in archival tissues whereas genes expressed at low levels in liver had no signal and did not vary from input control (Figure S5). In both fresh and archival laboratory mouse liver tissue, we observed significant MNase signal changes at 60-75% of genes with high expression (zFPKM > 2) compared to approximately 25% of genes with low expression (zFPKM < −2) (Figure S4A). Unsurprisingly, given the FAIRE assay’s low signal-to-noise ratio, signal changes showed a relatively weaker and more variable association with gene expression with changes observed at 25-80% of genes with high expression (zFPKM > 2) compared to 0-20% of genes with low expression (zFPKM < −2) (Figure S4B). Observation of relative signal strengths of the MNase and FAIRE assays in archival tissues consistent with expectations for their relative performance fresh tissues provides compelling evidence for characterising archival chromatin. The versatility of multiple approaches also suggests the feasibility of adapting other chromatin profiling assays, such as ChIP-Seq, Hi-C and ATAC-Seq (Assay for Transposase-Accessible Chromatin [19]), for use in archival tissues.

As the final demonstration of our novel approach, we characterised archival chromatin architecture in truly historical formaldehyde-preserved museum specimens obtained from the Queensland Museum. We selected five eastern water dragon (*Intellegama lesueurii lesueurii*) specimens preserved with formaldehyde between 1905 and 2001 (Table 2). Real museum specimens are a finite and precious resource, thus we only had liver tissue volume (29 – 200 mg) sufficient for a single archival chromatin assay. We selected archival MNase due to its stronger occupancy signal, superior repeatability, and semi-quantitative association with gene expression. We note that whilst FAIRE was not conducted on these samples, it is still a valuable tool for future independent verification of MNase results. Given that we expected to achieve relatively low coverage from whole genome sequencing of the archival tissues, we also sequenced a modern fresh tissue genomic DNA extraction from liver to serve as input to control for biases associated with the underlying sequence.

**Table 2.** Specimen details for archival eastern water dragon specimens. We selected five archival eastern water dragon (*Intellegama lesueurii lesueurii*) specimens for gDNA and MNase processing. For each specimen, we provide the Queensland Museum (QM) registration number, Habitat type, Latitude and Longitude of the collection location, the Time Interval in years between preparation & fixed-tissue sampling, Sex of the individual as well as pH and residual formaldehyde concentration ([F] as mg/L) of the specimen media at the time of sampling. Note, all collection localities are in Queensland, Australia.

| Sample name | QM Reg. No. | Habitat | Latitude | Longitude | Time Interval (years) | Sex | pH | [F] |
| --- | --- | --- | --- | --- | --- | --- | --- | --- |
| 1905M | J91140 | Urban | -27.46666667 | 153.0166667 | 116 | M | 6.78 | 400 |
| 1990M | J51722 | Urban | -27.35 | 152.9666667 | 31 | M | 6.53 | 200 |
| 1992M | J54438 | Non-urban | -27.91666667 | 152.3333333 | 29 | M | 6.53 | 200 |
| 2001F | J76769 | Non-urban | -27.83333333 | 153.1666667 | 20 | F | 6.53 | 200 |
| 2001M | J76089 | Urban | -27.51666667 | 152.95 | 20 | M | 6.53 | 200 |

All five water dragon samples produced clear evidence of genome-wide occupancy signal changes in regions ±2 kb of TSSs (Figure S6). These are the first ever historical chromatin accessibility profiles for fixed soft tissues, now validated in specimens up to 112 years old. Even with the limited sample sizes in this study, our experimental mouse data clearly demonstrates the feasibility of inferring historical gene regulation. For both vertebrate systems, the predominant occupancy signal is associated with phenotypic sex (Figure 5A & C). Males and females cluster along the first PC axis using a Pearson correlation analysis and explain a large proportion of the variation in chromatin profiles (Figure 5A; Mouse PC1 = 97.8%; Water dragons = 76.5%). Interestingly, the magnitude of correlation within the water dragon analyses is roughly 20-fold that of those within the mouse analyses, perhaps indicating that sex has a stronger influence on genome-wide MNase signal in water dragons which could reflect the greater influence of epigenetic processes over sexual phenotype in species with environmental sex determination [20]. Alternatively, it could be consistent with the occurrence of somatic cell-autonomous sex-identity, a phenomenon previously observed in some birds and reptiles [21–24].

**Figure 5.**
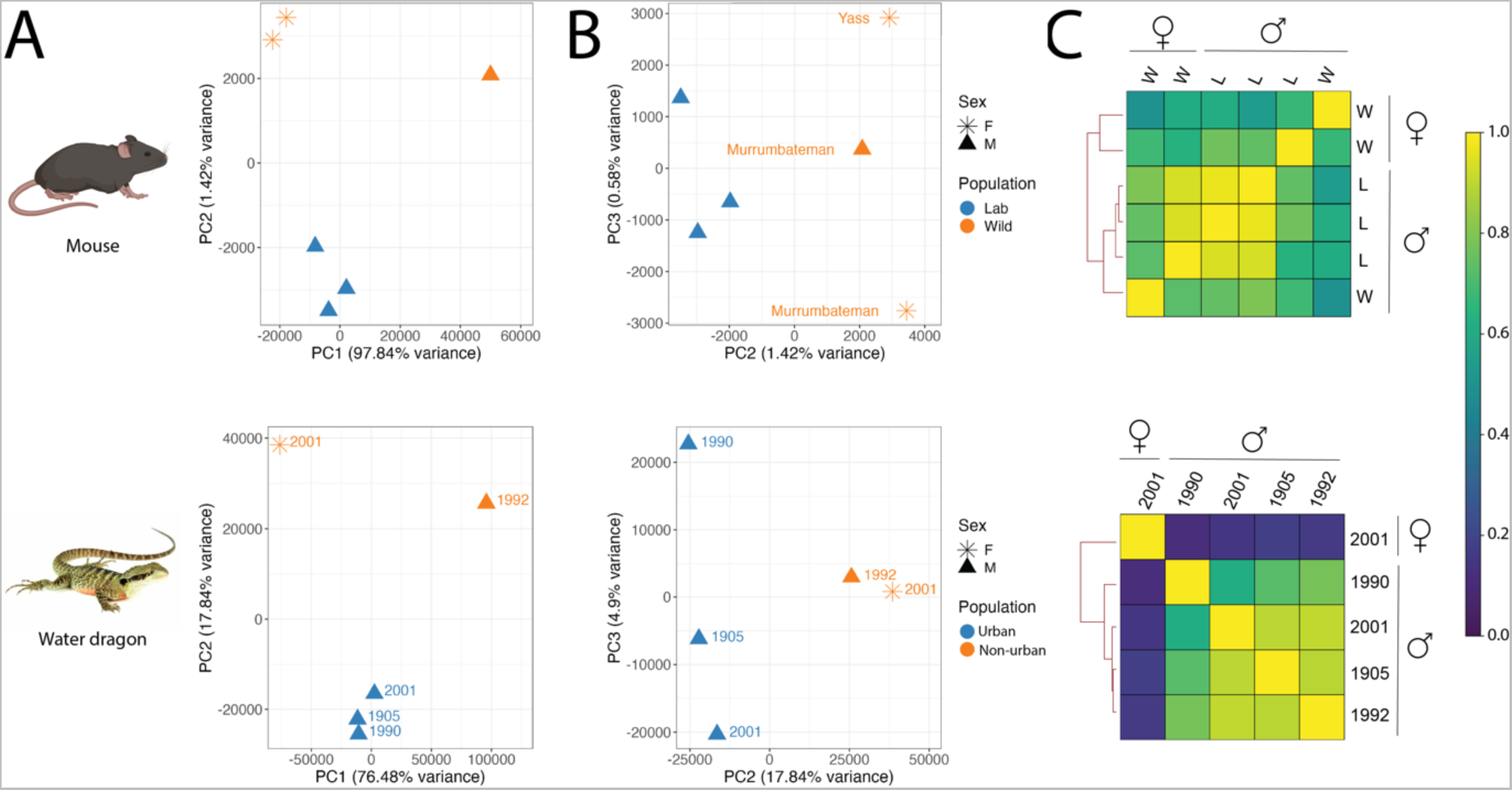
Sex and population signatures in heavily fixed archival specimens. In two species (mouse and water dragon), Pearson correlation-based analysis of genome-wide archival MNase occupancy signals resolves clusters of individuals by sex and population. PCA plots of principal components one and two (**A**) illustrate strong separation of females from males while plotting components two and three (**B**) reveals clustering of individuals along PC2 in accordance with population. For mouse, only the autosomal chromosomes were considered for this analysis. The shape of individual PCA plot points indicates the specimen’s sex (F = star; M = triangle) and color indicates the population (mouse – blue = laboratory, orange = wild; water dragon – blue = urban, orange = non-urban). Wild mice are labelled by collation location and water dragons are labelled by collection date. (**C**) Representing the Pearson correlation analysis as a heatmap, a relatively stronger differentiation of the sole female water dragon from the four males emerges in comparison to the sex-based differentiation observed in mouse. In the mouse heatmap, as in the PCA plots, the individuals cluster first by sex and then by population (L = lab; W = wild).

Eliminating the strong influence of sex in PC1 and instead comparing PC2 and PC3, both mouse and water dragon chromatin profiles segregate into groups consistent with habitat type at the time of collection (Figure 5B). Specifically, water dragons collected in urban Brisbane cluster separately to individuals from non-urban bushland habitats; and lab mice form a tight, highly reproducible cluster quite distinct from the more genetically and transcriptionally diverse outbred wild mice. Increased sampling will allow investigation of the degree to which this signal is influenced by genetic similarity and population structuring. In this small sample set, we see an indication that genetic similarity alone is not the sole driver of MNase clustering in that the three wild mice separate from the laboratory mice but do not themselves cluster by sampling location. Our findings also underscore the importance of sex-matching when selecting individuals for future work aiming to measure environmental effects.

Lastly, our water dragon results recapitulate previous results from our group that show that specimen age is a poor predictor of sequencing suitability [10]. Here, the 1905 specimen represents the oldest formaldehyde-preserved museum specimen to be successfully sequenced to date and the MNase reads alone yielded the highest whole genome cover yet achieved in archival formaldehyde-preserved specimens (genome cover = 5-8X; Table 3). Thus, our archival MNase method is state-of-the-art for obtaining both genomic and epigenomic data from formaldehyde-preserved specimens and does so simultaneously.

**Table 3.** Extraction and sequencing details for vertebrate specimens. Summary of the extraction and sequencing results for six mouse specimens (both fresh and archival tissues) and five eastern water dragon specimens (archival only) processed with archival MNase treatment. For each specimen, we processed the available mass of archival liver tissue (reported in mg) and report DNA yield in ng/mg. For each mouse specimen, we processed a 30 mg section of fresh tissue. For all tissues, we report DNA yield in ng/mg of tissue, the number of raw read pairs (millions), the percentage of raw reads mapping to the reference genome, the mean insert size of mapped reads in base pairs and the mean genome coverage after de-duplication and GC correction.

|  | Archival tissue weight (mg) | DNA yield (ng/mg tissue) |  | Raw read pairs (M) |  | Reads mapping (%) |  | Mean insert size (bp) |  | Mean genome cover (X) |  |
| --- | --- | --- | --- | --- | --- | --- | --- | --- | --- | --- | --- |
| *Mouse specimens* |  | Fresh | Archival | Fresh | Archival | Fresh | Archival | Fresh | Archival | Fresh | Archival |
| Lab1 | 287 | 28 | 14.4 | 282 | 250 | 82 | 67 | 178 | 129 | 29.2 | 15.3 |
| Lab2 | 545 | 52 | 7.8 | 293 | 199 | 81 | 68 | 101 | 124 | 16.9 | 11.9 |
| Lab3 | 593 | 45 | 7.6 | 249 | 228 | 83 | 65 | 97 | 124 | 14.2 | 13.1 |
| Wild1 | 467 | 44 | 4.5 | 306 | 239 | 84 | 67 | 115 | 117 | 21.1 | 13.3 |
| Wild2 | 624 | 21 | 6.8 | 262 | 263 | 93 | 67 | 124 | 118 | 21.4 | 14.9 |
| Wild3 | 763 | 77 | 3.5 | 256 | 227 | 85 | 68 | 128 | 112 | 19.7 | 12.2 |
| *Water dragon specimens* |  |  |  |  |  |  |  |  |  |  |  |
| 1905M | 82 | 0.08 |  | 146 |  | 40 |  | 74 |  | 5 |  |
| 1990M | 29 | 0.24 |  | 151 |  | 28 |  | 65 |  | 3.2 |  |
| 1992M | 45 | 0.12 |  | 171 |  | 32 |  | 68 |  | 4.3 |  |
| 2001F | 200 | 0.25 |  | 162 |  | 49 |  | 76 |  | 7.1 |  |
| 2001M | 92 | 0.63 |  | 154 |  | 58 |  | 78 |  | 8.1 |  |

## Discussion

Our new perspective on the utility of formaldehyde-fixed archival specimens provides the first multi-organ historical epigenetic capability. Our methods open the door to systematic and comprehensive investigation into the temporal dynamics of chromatin accessibility by drawing upon the untapped potential of museums and global biorepositories. Contrary to the prevailing dogma, we have shown that over-fixation with formaldehyde does not destroy DNA but rather enables successful recovery of historical chromatin architecture. This new capability has the potential to revolutionise the power of modern epigenome-wide association studies, in the pursuit of functional regulatory variants, by characterising vertebrate chromatin architecture over the last century.

Broad adoption of archival chromatin techniques by the world’s natural history collections and their users will require careful sampling designs to control for the effects of post-mortem degradation, specimen sex, genetic background, and age. While the interval between death and fixation is rarely if ever recorded, our group as previously reported that the integrity of the gut contents can be used as a proxy for degradation when vetting specimens [10]. Sex-specific gene expression is observed across a wide range of vertebrate tissues, even those that are not gonadal in origin or associated with secondary sexual characters [25]. Thus, controlling for sex should be a key consideration in any study design. Likewise, age of the individuals should be considered given expected changes in chromatin accessibility associated with aging [26,27]. We observed greater MNase signal variation in the wild mouse samples compared to laboratory mice likely due to a combination of factors, such as variation in sex, age, diet, exercise, or genetic background. This indicates that a higher degree of replication will be required within carefully matched specimen sets to study historical wild populations.

Now that recovery of genomic data from formaldehyde-fixed museum specimens has been firmly established by this and other studies, we should reassess our assumptions about the damaging effect of fixation on other nucleic acids and epigenetic modifications. For example, could quantitatively informative mRNA feasibly be retrieved from historical specimens? Precedents have been set by successful research on clinical FFPE samples [28], and a single study has recently retrieved RNA from formaldehyde-fixed museum specimens [29]. A better understanding of how fixation modulates the molecular signal in these contexts could revolutionise our ability to study long-term temporal trends in gene regulation. We hope that the methods described here are the first of a suite of approaches to characterise historical gene regulation. Achieving this feat will increase the power of modern studies seeking to imply causal functional relationships between regulatory variation and phenotypes, just as the baseline data produced by ancient DNA genomic analysis has accelerated the identification of functional human sequence variation [30].

## Methods

### Yeast processing

We used established procedures for MNase [13] and FAIRE [12] with yeast and made optimisations for processing heavily fixed cultures.

#### Culture, fixation, and preparation of nuclei

We inoculated 500 mL YPD media from an overnight culture of *Saccharomyces cerevisiae* strain BJ5464 auxotroph ΔURA3 and grew to OD600 = 0.75 with shaking at 28°C. We split the remaining culture into two equal volumes and incubated one flask under heat stress conditions in a 37°C water bath with gentle shaking for 20 minutes. At this pre-fixation stage, we removed 2 mL aliquots from both the optimal growth and heat shock flasks for gDNA and RNA extraction.

To both the optimal growth and heat shock flasks, we added formaldehyde to a final concentration of 1% and began incubation with slow shaking at room temperature. At 15 min, 1 hr, 6 hr and 24 hr we collected aliquots and quenched fixation with addition of glycine to a final concentration of 0.125 M followed by gentle shaking at room temperature for 15 min. We pelleted the fixed cells by centrifugation at 4,000 × *g* for 5 min at 4°C and washed twice with 15 mL of cold Phosphate Buffered Saline, pH 7.4 (PBS). To fully quench any remaining formaldehyde, we resuspended the pellets in 10 mL cold Glycine Tris EDTA (GTE; 100 mM glycine, 10 mM Tris-HCl, pH 8.0, 1 mM EDTA) buffer and incubated them at 4°C with rocking for 24 hr.

To isolate nuclei from the fixed cells, we harvested the cells by centrifugation at 4,000 × *g* for 5 min at 4°C and washed twice with cold milliQ water before transferring the cells to 2 mL tubes. We resuspended the cell pellets in 2 mL Spheroplast Buffer (SB; 1M Sorbitol, 50 mM Tris, pH 7.5 with freshly added 10 mM β-mercaptoethanol), added Zymolase 20T (MP Biomedicals) to a final concentration of 0.25 mg/mL and incubated at 28°C for 2 hr. We harvested the fixed spheroplasts by centrifugation at 1,500 × *g* for 10 min at 4°C and washed twice with cold SB, resuspending in 10 mL. We then aliquoted the spheroplasts such that each aliquot contained a volume of cells from 4 mL and 10 mL of original culture for the short (15 min & 1 hr) and long (6 hr and 24 hr) cultures, respectively. We collected a larger volume of cells from the longer fixation timepoints to account for a lower expected DNA recovery due to heavy fixation. We pelleted the aliquoted spheroplasts once more at 1,500 × *g* for 10 min at 4°C, removed the supernatant and froze the tubes at −80°C until further processing.

#### gDNA and RNA extraction from unfixed yeast

We extracted gDNA and RNA from aliquoted unfixed cells frozen at −80°C. For gDNA extractions, we resuspended cells in 200 μL SB, added Zymolase 20T to a final concentration of 0.25 mg/mL and 1 μL RNase A and incubated at 37°C for 30 min. We harvested the spheroplasts by centrifugation at 3,000 × *g* for 10 min then resuspended the pellet in 100 μL PBS plus 0.01 M EDTA, added 2 μL of proteinase K (20 mg/mL) and incubated at 56°C for 45 min in a thermal mixer with agitation at full speed (1400 rpm). We purified gDNA from lysates with one phenol:chloroform:isoamyl alcohol (25:24:1) extraction and concentrated the gDNA on beads as described in *<u>Small fragment-optimised bead purification of DNA</u>*, eluting in 20 μL 10 mM Tris, pH 8.0. For RNA extractions, we resuspended three aliquots per culture in 450 μL RLT Buffer and followed the manufacturer’s instructions for the Qiagen RNeasy plant mini kit, eluting in 30 μL nuclease-free water.

#### Yeast MNase treatment

We resuspended triplicate aliquots of each fixation timepoint for both heat shock and optimal growth conditions in 200 μL MNase Digestion Buffer (DB; 0.5 mM spermidine, 0.075% Nonidet P40, 50 mM NaCl, 10 mM Tris pH 8.0, 5 mM MgCl_2_, 5 mM CaCl_2_ plus freshly added cOmplete EDTA-free protease inhibitor cocktail (Merck)), added 0.5 U of micrococcal nuclease (Worthington Biochemical Corporation) and incubated the tubes for 7 min in a 37°C water bath. After incubation, we immediately stopped the reaction with the addition of 50 μL of Quenching Solution (QS; 4% Triton X-100, 1.2% SDS, 600 mM NaCl, 12 mM EDTA) and incubated the tubes on ice for 10 min. We collected the digested chromatin by centrifuging at 14,000 × *g* for 10 min at 4°C and removing the supernatant to a new tube. We digested the supernatant with proteinase K at a final concentration of 0.1 mg/mL and incubated at 56°C for 2 hr. We further purified the DNA with two phenol:chloroform:isoamyl alcohol (25:24:1) extractions, treating with RNase (1 μL RNase A and incubation at room temperature for 30 min) between extractions one and two followed by *<u>Small fragment-optimised bead purification</u> <u>of DNA</u>*, resuspending in 20 μL 10 mM Tris, pH 8.0. We fully de-crosslinked the DNA with incubation at 65°C overnight. <u>Yeast FAIRE treatment</u>

We resuspended four aliquots of each fixation timepoint for both heat shock and optimal growth conditions in 1 mL Chromatin Shearing Buffer (CSB; 10 mM Tris-HCl pH 8.0, 0.1% SDS, 1 mM EDTA) and transferred each 1 mL suspension to a 1 mL Covaris milliTUBE. We sonicated the tubes in a Covaris E220 focused-ultrasonicator on settings PIP 420, duty factor 30%, cycles per burst 200 for 7 min (short fixation time points) or 12 min (long fixation time points). We transferred each aliquot of sonicated nuclei to a 2 mL tube and clarified the lysate by centrifugation at 5,500 × *g* for 15 min at 4°C. We set aside one tube per sample type at this point as an input control. With the remaining three tubes per sample type, we performed a phenol:chloroform:isoamyl alcohol (25:24:1) extraction including back-extraction of the organic phase with addition of 150 μL 10 mM Tris, pH 8.0 followed by an additional phenol:chloroform:isoamyl alcohol extraction. To the aqueous phase recovered from these tubes as well as the input control tubes set aside earlier, we added 10 μL RNase A and incubated at room temperature for 30 min followed by addition of 2 μL proteinase K and incubation at 55°C for 1 hr. We de-crosslinked overnight with incubation at 65°C and concentrated the DNA *<u>Small fragment-optimised bead purification of DNA</u>*, resuspending in 20 μL 10 mM Tris, pH 8.0.

### Mock archival specimen preparation

To thoroughly test archival chromatin methods in a species with a well-annotated genome, we created a bank of experimental formaldehyde-preserved in-bred laboratory and out-bred wild-trapped mouse specimens. We acquired male mice (*Mus musculus* strain C57BL/6) aged 17-18 weeks from Australian BioResources and sacrificed them upon arrival by cervical dislocation (Australian Ethics Committee number 2017-34). The CSIRO Health and Biosecurity Rodent Management Team donated adult *M. musculus* live trapped at two locations in May of 2019 (Murrumbateman, NSW, Australia, lat. −35.0424064, long. 148.99947; Yass, NSW, Australia, lat. −34.8682227, long. 149.00763) (Australian Ethics Committee number 2018-46). The wild mice had been housed according to standard mouse husbandry practices for 3 months prior to sacrificing them by cervical dislocation in August of 2019.

In accordance with modern archival procedure, we sampled liver tissue from all specimens for storage at −80°C to serve as a source of specimen-matched fresh tissue. We then prepared each carcass by emersion in 10% neutral buffered formalin (3.7% formaldehyde) for 3 days followed by soaking in water for 1 day before transfer to 70% ethanol. We archived these formaldehyde-preserved mice (hereafter referred to as archival mice) in the Australian National Wildlife Collection (ANWC; Crace, ACT, Australia) spirit vault in glass specimen jars.

### Vertebrate specimen selection and archival tissue sampling

#### Archival mice

We dissected archival liver tissue from our mock specimen collection housed at the ANWC and transferred the tissue to ethanol filled tubes for transport and further processing. At the time of dissection, the formaldehyde-fixed laboratory and wild-caught mice had been archived for 4.8 and 2.3 years, respectively.

#### Archival eastern water dragons

Prior to sampling archival eastern water dragons (*Intellegama lesueurii lesueurii*), we assessed the sequencing suitability of specimens archived at the Queensland Museum. Following established methods [10], we took aliquots of the preservation media and measured pH using an Orion Versa Star Pro benchtop pH meter (Thermo Scientific) and residual formaldehyde concentration ([*F*]) using MQuant test strips (Merck). From visually well-preserved specimens within jars registering neutral pH (6 > pH < 8) and low formaldehyde ([F] < 10,000 mg/L) we sampled archival liver tissue and transferred the tissue to ethanol-filled tubes for transport and further processing.

### Archival vertebrate tissue processing

The following describes the final optimised tissue processing procedure we used on all mouse and eastern water dragon specimens.

#### Tissue preparation

We pulverized and rehydrated archival tissues similarly as in [10]. We cryo-pulverised the tissues into a rough powder using a cryoPREP (Covaris) automated dry pulverizer (3 impacts to an extra thick TT1 TissueTube on intensity setting 6 with immersion in liquid N_2_ for 10 seconds between impacts). We rehydrated the pulverized tissue under ice-cold conditions by stepping into 50% ethanol, 30% ethanol then water with rocking at 4°C for 10 min intervals with collection of the tissue by centrifugation at 4000 × *g* for 5 min at 4°C. We quenched residual formaldehyde by rocking overnight at 4°C in an excess volume (approximately 15 mL to 50 mg tissue) GTE buffer.

#### Isolation of archival nuclei

We centrifuged prepared tissue for each specimen at 4000 × *g* for 5 min at 4°C and washed once with ice-cold phosphate buffered saline (PBS). As an initial step to improve tissue dissociation, we resuspended the tissue in 1 mL of pre-heated Pepsin solution (0.5% Pepsin in 5 mM HCl, pH 1.5) then incubated the tubes at 37°C in a ThermoMixer (Eppendorf) for 90 min with rotation set at 750 rpm. We then performed three washes with cold PBS and resuspended in 1 mL sodium citrate buffer (pH 6) before transferring the suspension to a 2 mL glass Dounce homogenizer. For fine tissue dissociation, we homogenized the tissue on ice until the larger chucks were broken up and pestle moved freely (20-30 strokes). To improve nuclei isolation, we performed initial de-crosslinking by transferring the tissue to a 2 mL tube and incubating at 80°C in the sodium citrate buffer for 1 hr. We then washed the tissue three times with ice-cold PBS before resuspending in 1 mL ice-cold Farnham Lysis Buffer (FLB; 5 mM PIPES pH 8.0, 0.1% SDS, 1 mM EDTA) and transferring to a 1mL Covaris milliTUBE containing an AFA fibre. We isolated nuclei by NEXSON [31] with sonication of the tubes in a Covaris E220 focused-ultrasonicator on settings PIP 160, DF 15%, CBP 200 for 600 seconds. With the mouse samples, we split the tissue in half to process with MNase and FAIRE. With all tubes, we pelleted the nuclei, removed the supernatant, and froze the pellets at –80°C.

#### Archival MNase treatment

We based the archival MNase protocol on [18] with substantial optimisation for heavy fixed input. We resuspended the frozen nuclei in 200 μL MNase DB per 50 mg tissue. To each tube we added 0.5 U MNase (Worthington Biochemical Corporation) and 200 U Exonuclease III (New England Biolabs) per 50 mg of tissue and incubated at 37°C with 750 rpm rotation for 15 min. To quench the digestion, we immediately added 50 μL QS per 50 mg of tissue, mixed well and incubated on ice for 10 min. To enhance release of the digested chromatin from the nuclear debris, we transferred the suspension to a 1 mL Covaris miliTUBEs containing an AFA fibre and briefly sonicated the samples in a Covaris E220 focused-ultrasonicator with settings PIP 160, DF 15%, CBP 200 for 60 seconds. We transferred the sonicated digest to a new 2 mL tube and clarified by centrifugation for 10 min, 9600 × *g*, at 4°C, transferring the supernatant to a new tube. We added 1 μL RNaseA and incubated for 30 min at room temperature followed by addition of 2 μL 20 mg/mL proteinase K and incubation at 55°C for 1 hour. We purified DNA fragments with a phenol:chloroform:isoamyl alcohol (25:24:1) extraction including back-extraction of the organic phase with addition of 150 μL 10 mM Tris, pH 8.0 followed by an additional phenol:chloroform:isoamyl alcohol extraction. We de-crosslinked overnight with incubation at 65°C and concentrated the DNA via *<u>Small fragment-optimised bead purification</u> <u>of DNA</u>*, resuspending in 20 μL 10 mM Tris, pH 8.0.

#### Archival FAIRE treatment

We based the archival FAIRE protocol on [12,32] with modifications to chromatin shearing and extractions. We resuspended nuclei in 1 mL chromatin shearing buffer (10 mM Tris-HCl pH 8.0, 0.1% SDS, 1 mM EDTA) and transferred the suspension to new 1 mL Covaris milliTUBEs containing an AFA fibre. We sheared the chromatin via sonication in a Covaris E220 focused-ultrasonicator with settings PIP 420, DF 30%, CBP 200 for 10 min. We clarified the lysate by centrifugation for 15 min 5,500 × *g* at 4°C, removed the supernatant to a new tube. We added 1 μL RNaseA and incubated for 30 min at room temperature. At this point, we reserved 10% of the sheared chromatin to purify as an input control. With the FAIRE fraction, we depleted protein-bound DNA through extraction with phenol:chloroform:isoamyl alcohol including back-extraction of the organic phase with addition of 150 μL 10 mM Tris, pH 8.0 followed by an additional phenol:chloroform:isoamyl alcohol extraction. To the reserved input control, we added 2 μL proteinase K solution and incubated at 65°C for 1 hour and de-crosslinked both the FAIRE and input fractions overnight with incubation at 65°C. We performed two rounds of phenol:chloroform:isoamyl alcohol extraction with back extraction upon the input controls and concentrated FAIRE and input fraction DNA via *<u>Small fragment-optimised bead purification of DNA</u>*, resuspending in 20 μL 10 mM Tris, pH 8.0.

### Fresh Vertebrate Tissue Processing

#### Fixation of fresh tissue

On dry ice, we transferred approximately 30 mg of liver tissue from each flash-frozen specimen stock to an extra thick TT05 TissueTube and cryo-pulverised the tissues into a rough powder using a cryoPREP (Covaris) automated dry pulverizer (2 impacts on intensity setting 6 with immersion in liquid N_2_ for 10 seconds between impacts). We resuspended the pulverised tissue in 1.5 mL room temperate PBS, transferred the suspension to a 2 mL tube and immediately added 40 μL 37% formaldehyde for a final concentration of 1% formaldehyde. We rocked the tube at room temperature for 15 min and then quenched fixation through the addition of 79 μL 2.5 M glycine to achieve a final concentration of 125 mM glycine. We continued to rock the suspension for 5 min at room temperature and then centrifuged 500 × *g* for 5 min at 4°C.

#### Isolation of nuclei from fixed frozen tissue

We washed the pulverised fixed tissue three times with ice-cold PBS, resuspended in 1 mL FLB and transferred the tissue to a 1 mL Covaris milliTUBE containing an AFA fibre. We isolated nuclei by NEXSON in a Covaris E220 focused-ultrasonicator with settings PIP 150, DF 10%, CBP 200 for 300 seconds. We split the tissue in half to process with MNase and FAIRE. With all tubes, we pelleted the nuclei, removed the supernatant, and froze the pellets at –80°C.

#### Fixed frozen tissue MNase treatment

For MNase treatment of the fixed frozen mouse tissues, we adapted [18] to conform with the equipment used in our modified archival MNase protocol. To the pelleted nuclei, we added 200 μL DB and gently resuspended. To each tube we added 0.5 U MNase (Worthington Biochemical Corporation) and 200 U Exonuclease III (New England Biolabs) and incubated at 37°C with 750 rpm rotation for 15 min. To quench the digestion, we immediately added 50 μL QS, mixed well and incubated on ice for 10 min. To enhance release of the digested chromatin from the nuclear debris, we transferred the suspension to a 1 mL Covaris miliTUBEs containing an AFA fibre and briefly sonicated the samples in a Covaris E220 focused-ultrasonicator with settings PIP 160, DF 15%, CBP 200 for 60 seconds. We transferred the sonicated digest to a new 2 mL tube and clarified by centrifugation for 10 min, 9600 × *g*, at 4 °C, transferring the supernatant to a new tube. We added 1 μL RNaseA and incubated for 30 min at room temperature followed by addition of 2 μL 20 mg/mL proteinase K and incubation at 55°C for 1 hour. We purified DNA fragments with a phenol:chloroform:isoamyl alcohol (25:24:1) extraction including back-extraction of the organic phase with addition of 150 μL 10 mM Tris, pH 8.0 followed by an additional phenol:chloroform:isoamyl alcohol extraction. We de-crosslinked overnight with incubation at 65°C and concentrated the DNA via *<u>Small fragment-optimised bead purification of DNA</u>*, resuspending in 20 μL 10 mM Tris, pH 8.0.

#### Fixed frozen tissue FAIRE treatment

We followed an established FAIRE protocol [12,32] for processing fresh tissues with modifications to conform with the equipment used in our archival FAIRE protocol. To the pelleted nuclei, we added 1 mL chromatin shearing buffer (10 mM Tris-HCl pH 8.0, 0.1% SDS, 1 mM EDTA), resuspended and transferred the suspension to new 1 mL Covaris milliTUBEs containing an AFA fibre. We sheared the chromatin via sonication in a Covaris E220 focused-ultrasonicator with settings PIP 420, DF 30%, CBP 200 for 12 min. We clarified the lysate by centrifugation for 15 min 5,500 × *g* at 4°C and removed the supernatant to a new tube. We added 1 μL RNaseA and incubated for 30 min at room temperature. At this point, we reserved 10% of the sheared chromatin to purify as an input control and further processed the FAIRE and input controls as we did the archival samples.

#### RNA extraction from fresh mouse tissues

We extracted RNA from fresh mouse using an AllPrep DNA/RNA kit (Qiagen). We placed 5-10 mg of tissue into 350 μL RLT Plus Buffer in a 2 mL tube containing a 5mm stainless steel bead and homogenized with a TissueLyzer for two 2 min rounds at 30 Hz. We proceeded to follow manufacturer’s instructions to isolate RNA, eluting in 30 μL RNase-free water.

#### Preparation of water dragon input control

The five archival specimens lacked sufficient archival tissue for processing of an input control and were not archived with specimen-matched fresh tissue. Thus, we opted to procure fresh tissue from three separate individuals to serve as a pooled input control. We dissected frozen liver tissue from animal which had been euthanised due to injury in accordance with Queensland Department of Environment and Sciences permit WA0038029 (Australian Ethics Committee number ANA20161, University of Sunshine Coast). Using a TissueLyzer, we pulverized approximately 5 mg of tissue per specimen in a 2 mL tube containing a 5mm stainless steel bead. We then immediately added 350 μL RLT Plus Buffer and followed manufacturer’s instructions for the Qiagen AllPrep kit, eluting in 100 μL Elution Buffer.

#### Small fragment-optimised bead purification of DNA

We concentrated purified DNA with a custom small-fragment optimised SPRI bead clean-up procedure. Our bead solution is prepared in 50 mL aliquots from 1 mL Sera-Mag (Cytiva) beads such that when added to the DNA extract in a ratio of 1.5:1(bead solution:DNA) the final concentration of reagents equal 12% PEG-8000, 40% isopropanol, 0.6 M NaCl, 6 mM Tris-HCl (pH 8.0), 0.6 mM EDTA and 0.03% Tween-20. After adding the bead solution, we incubated the tubes at room temperature for 15 min with rotation and then pelleted the beads on a magnet and removed the supernatant. We then resuspended the beads in fresh 70% ethanol, pelleted them upon a magnet and washed the beads once more with 70% ethanol. After removing the ethanol and allowing the beads to dry briefly for 30 sec, we resuspended the beads in 20 μL 10 mM Tris EDTA and incubated the tubes at 37°C for 15 min to thoroughly elute the DNA. Finally, we pelleted the beads on a magnet and transferred the supernatant to a new tube.

### Nucleic acid quantification

We quantified DNA by Qubit (1X dsDNA HS Assay kit) and Tapestation (High Sensitivity D1000), following the manufacturer’s instructions. We quantified the RNA by NanoDrop and Bioanalyzer (Total RNA Nano), following the manufacturer’s instructions. We report the DNA yield from the vertebrate tissue samples in Table 3.

### Library Preparation & Sequencing

The Australian Genome Research Facility (AGRF) performed all library preparation and sequencing. AGRF prepared all DNA libraries with the xGen cfDNA & FFPE DNA Library Prep Kit (IDT) and sequenced the yeast DNA libraries on a single 200 cycle (100 bp PE) Illumina NovaSeq S4 lane, all mouse and water dragon DNA libraries across four 300 cycle (150 bp PE) Illumina NovaSeq S4 lanes. AGRF prepared Illumina stranded mRNA libraries from the mouse RNA extracts and sequenced the pool on a 100 cycle (100 bp SE) NovaSeq S4 lane.

### Analyses

#### Genome preparation

For yeast we used the *S. cerevisiae* S288C R64/sacCer3 RefSeq assembly. For the *M. musculus* reference genome, we used the GRCm38.p6 RefSeq assembly. For *I. l. lesueurii,* we used the *Pogona vitticeps* pvi1.1 RefSeq assembly as a species-specific reference was not available. We masked repeats in all genomes with RepeatMasker v.4.1.0 (http://www.repeatmasker.org) through two rounds of masking – first with default settings and then again providing a list of standard Illumina adapters with -e rmblast enabled.

#### DNA read alignment

We computed quality control metrics for the raw reads using FastQC version 0.11.8 [33]. To facilitate deduplication with the IDT library unique molecular identifiers (UMIs), we converted the Fastq files to BAM format with FastqToSam in PICARD v 2.9.2 [34], extracted the UMIs with ExtractUmisFromBam in FGBio v. 1.3.0 [35] and restored the files to Fastq format with SamToFastq in PICARD. We aligned raw reads with the kalign function of the ngskit4b tool suite version 200218 [36] with options -c25 -l25 -d50 -U4. We removed PCR and optical duplicates from the alignments using the MarkDuplicates function of PICARD enabling REMOVE_DUPLICATES=TRUE and utilising UMIs. We computed and corrected for GC-bias with deepTools version 3.5.1 [37] using effective genome sizes of 12,157,105 bp, 2,818,974,548 bp and 1,716,675,060 bp and for the sacCer3, GRCm38.p6 and pvi1.1 and genomes, respectively. We calculated the mean aligned insert length using the CollectInsertSizeMetrics function of PICARD and estimated nuclear genome coverage as the number of unique aligned GC-corrected reads multiplied by the mean insert length divided by unmasked genome size. We report the sequencing yield and mapping results of all vertebrate samples in Table 3. We visualized alignments in CLC Genomics Workbench 21 (Qiagen).

#### Peak analyses

We analysed regional sequence depth enrichment as occupancy values with the dtriple function in DANPOS3 [14]. We analysed all yeast and mouse alignments both as individual samples and as pools of three replicates. For profiling the effect of FAIRE and MNase treatment, we used a corresponding input control. For differential peak analyses, we ran pooled heat-shocked cultures versus pooled optimal growth conditions as input control for yeast and pooled laboratory strain versus pooled wild caught as input for mouse. We analysed all water dragon alignments individually compared to a pool of three input control alignments. With the output of the DANPOS dpeak function, we enforced a significance cut-off of FDR < 0.05 upon sites with local peak gains, local peak losses and log2fold-change in total peak signal. We used the R packages *ChIPseeker* [38,39], *GenomicFeatures* and *GenomicRanges* [40] to annotate sites with significant peak changes and restrict our downstream analyses to the 2 kb region upstream of transcription start sites (TSSs).

#### RNA-Seq analysis

We aligned RNA-Seq reads from yeast and mouse to their respective masked genomes with kalign and calculated FPKM using the Tuxedo pipeline [41]. We then z-transformed the FPKM values using the R package *zFPKM* and created lists of genes with low (zFPKM < −2) medium (−2 > zFPKM < 2) and high (zFPKM > 2) expression [42]. To account for regions with poor mapping, we eliminated genes from the low score list with Z scores less than −20 (indicating reads mapped to the gene but at very low levels). For the differential expression analyses, we used the Trinity edgeR pipeline [43,44] to calculate log2FC between treatments.

#### Downstream analyses of peak profiles

Where we summarised shared genes with significant peak changes between replicates, we used the R package *ggVennDiagram* to generate Venn Diagrams. We calculated Gene Ontology (GO) Biological Process enrichment for yeast within lists of genes with significant peak changes as well a list of differentially expressed genes (as determined by RNA-Seq) using Yeast EnrichR [16,17]. For mouse, we calculated tissue-specific enrichment within lists of genes with significant peak changes using TissueEnrich [45]. We generated genome wide TSS peak enrichment heatmaps with the tagHeatmap function in *ChIPseeker*. To measure the effect of gene expression on upstream peak changes, we calculated the proportion of genes with low, medium and high expression with significant peak changes in the 2kb upstream region for each replicate and plotted the proportions as violin plots.

#### Pearson correlation analyses

To compare genome-wide MNase profiles between individuals for mouse and water dragon, we converted each individuals’ DANPOS dpeak wig file from having compared the MNase profile to input control to bigWig format and compiled a summary matrix for each species using Deeptools [37]. We performed principal component analysis upon the resulting summary matrices with the *prcomp* function in base R and plotted principal components one through three using *ggplot2*.

## Acknowledgements

We thank Olly Berry and Andrew Young for their leadership within the Environomics Future Science Platform. Dan Powell for technical assistance. We thank the director of the Australian National Wildlife Collection, Leo Joseph, and the ANWC staff (specifically, Margaret Cawsey, Alex Drew, Tonya Haff, and Chris Wilson) for their contributions of curatorial expertise, metadata management and sampling assistance. We thank Wendy Ruscoe for acquiring wild Australian mice and Oliver Mead for supplying yeast cultures. We thank Patrick Couper at the Queensland Museum for assistance in sampling the Eastern water dragons. We thank Ondrej Hlinka and CSIRO IM&T Client Services for their assistance in utilising the CSIRO supercomputing system. We thank the Australian Genome Research Facility for sequencing advice. We thank Don Gardiner, Kerensa McElroy, Yijin Liew, Cheng-Soon Ong and the Environomics Epigenetics Discussion Group for valuable comments on study design and analysis. We would like to acknowledge the contribution of Bioplatforms Australia in the generation of data used in this publication. Bioplatforms Australia is enabled by NCRIS.

Funding for this study was provided by the Environomics CSIRO Future Science Platform (grants R-10011 and R-14486) awarded to CEH.

