## Supplementary Figures for "Century-old chromatin architecture preserved with formaldehyde"

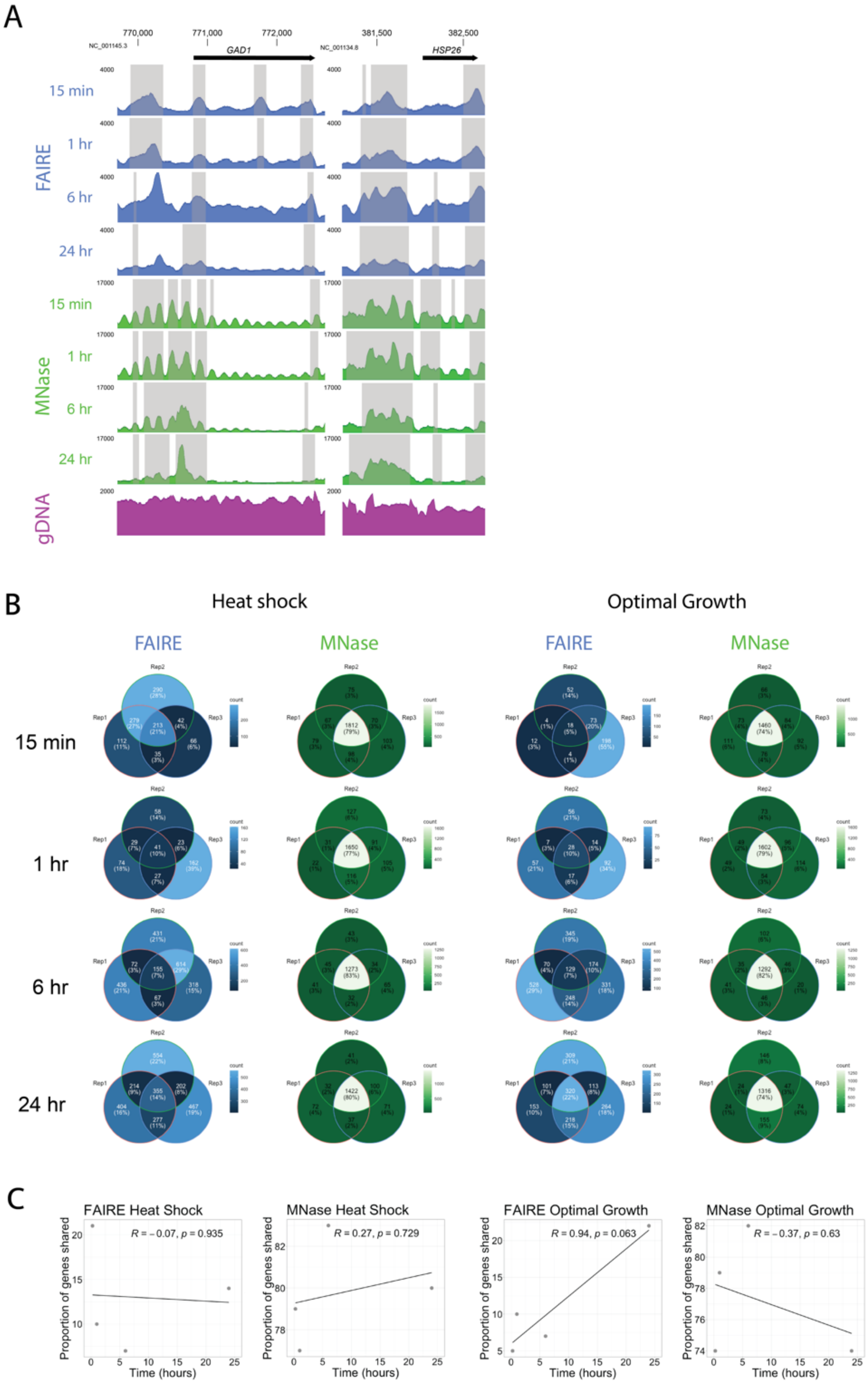

*Figure S1. Further Timepoints for Yeast alignments and Venn diagrams*

**A.** Pooled occupancy values (FAIRE: blue, MNase:green) compared to gDNA extraction control (purple) with formaldehyde fixation for 15 min, 1 hr, 6 hr and 24 hr of heat shocked *S. cerevisiae*. Shading indicates regions with significant peak width shifts (FDR < 0.05) between treatment and input control. Upstream of highly upregulated GAD1 and HSP26 genes (log2FC = 3.8 and 9.02), changes in occupancy signal morphology are observed. The 5' FAIRE peak broadens, while the distinct 5' MNase nucleosome array transforms into a single peak. **B.** Venn diagrams demonstrate repeatability of the FAIRE (blue) and MNase (green) assays among technical replicates in yeast cultures grown under heat shock or optimal growth conditions fixed with formaldehyde. Numbers/proportions represent genes with significant peak gain (FDR < 0.05, log10Pval < -6) within 2 kb upstream of the TSS. Lighter colors indicate higher shared gene count. **C.** Correlation tests between proportion of genes shared between all three replicates in (B) and fixation time. Linear regression lines are fitted, and correlation coefficients (R) and p-value are provided for each time point.

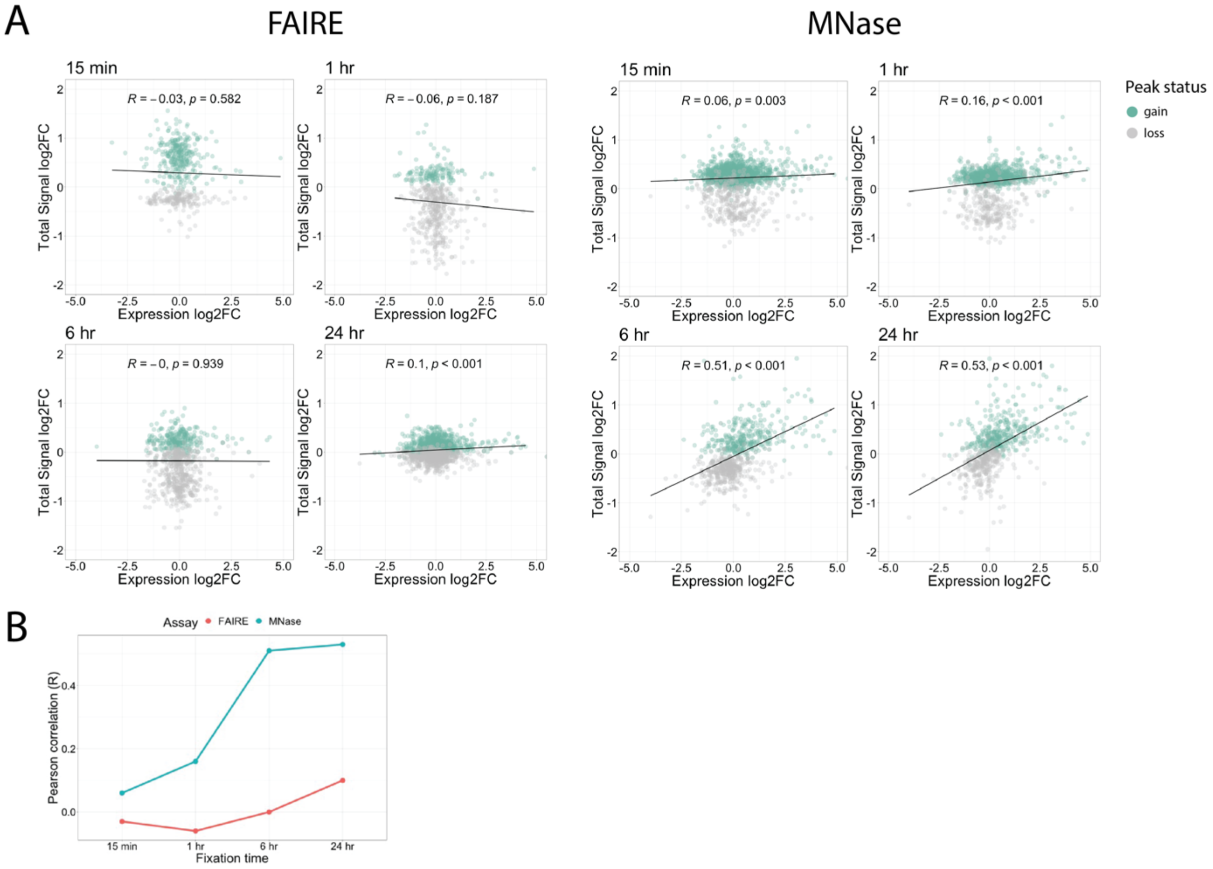


*Figure S2. Further Timepoints for yeast differential DANPOS & expression correlation*

**A.** For the FAIRE and MNase assay of yeast fixed for between 15 min and 24 hr, total signal log2FC for genes with significant (FDR < 0.05) total peak signal change between pooled replicate heat shock and optimal growth conditions in the 2 kb region upstream of the TSS is plotted against expression log2FC measured by RNA-Seq. Genes are coloured green for signal gain or grey for signal or loss. Linear regression lines are fitted, and correlation coefficients (R) and p-value are provided for each time point. **B.** Summary plot of Pearson correlation (R) values for the MNase and FAIRE time series in A.

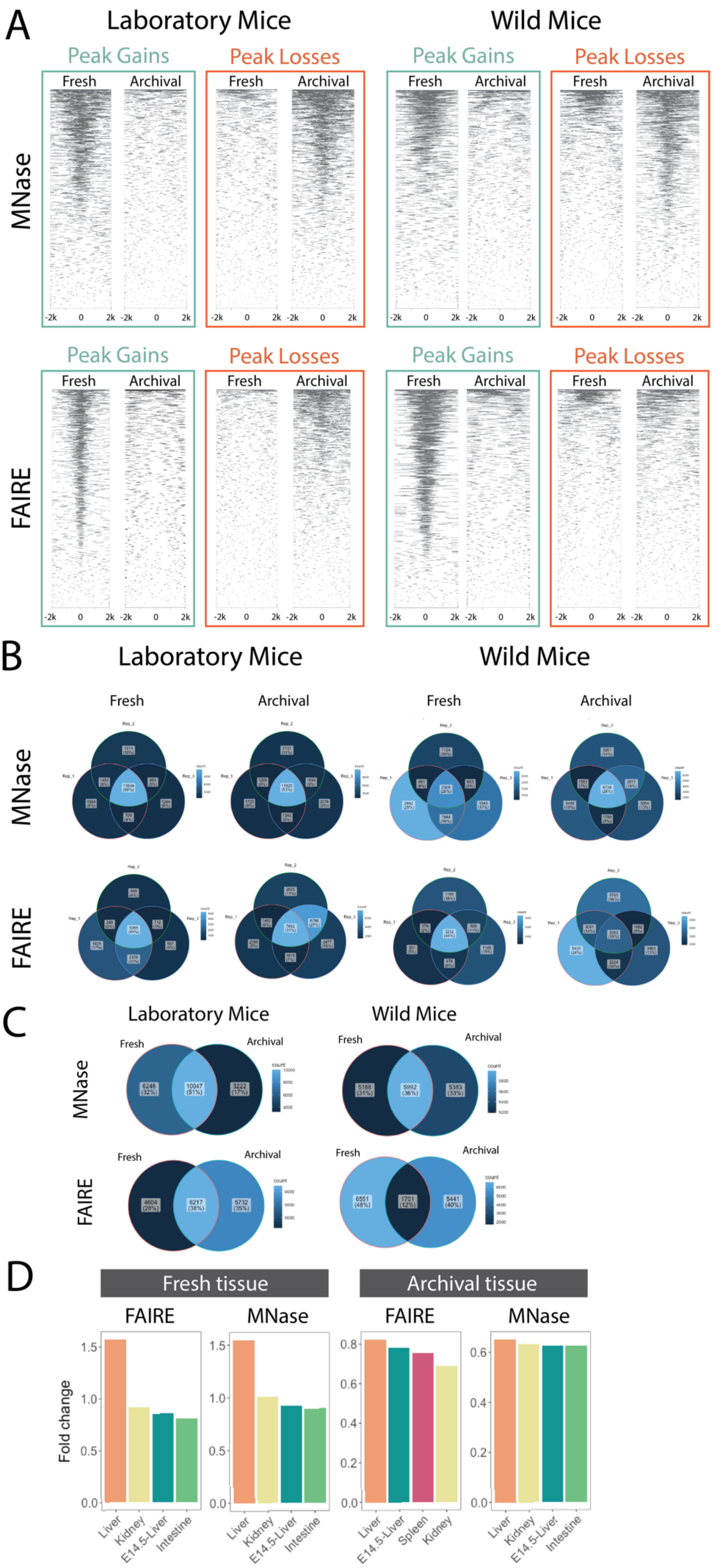

A
Laboratory Mice
Wild Mice
Peak Gains
Peak Losses
Fresh
Archival
MNase
FAIRE
-2k
0
2k
B
Laboratory Mice
Wild Mice
Fresh
Archival
MNase
FAIRE
C
Laboratory Mice
Wild Mice
Fresh
Archival
MNase
FAIRE
6248 (32%)
10047 (51%)
3222 (17%)
5168 (31%)
5992 (36%)
5383 (33%)
4604 (28%)
6217 (38%)
5732 (35%)
6551 (48%)
1701 (12%)
5441 (40%)
D
Fresh tissue
Archival tissue
FAIRE
MNase
Fold change
Liver
Kidney
E14.5-Liver
Intestine
Spleen

### *Figure S3. Further comparisons for mouse genome-wide signal*

**A.** Heatmap of MNase and FAIRE assay significant peak gains and losses (FDR < 0.05, log10Pval < -6) in fresh and archival tissues 2 kb either side of genome-wide transcription start sites pooled across three individuals of laboratory or wild mice. **B**. Venn diagrams demonstrating relative repeatability of the MNase and FAIRE assays applied to fresh and archival liver tissue among biological replicates in laboratory and wild mice. Numbers/proportions represent genes with significant peak gains for fresh tissue and losses for archival tissue (FDR < 0.05, log10Pval < -6) within 2 kb upstream of the TSS. Lighter colors indicate higher shared gene count. **C.** Venn diagrams showing overlap in genes identified through both fresh and archival MNase and FAIRE assays applied to three replicates in laboratory and wild mice. **D.** Tissue enrichment within shared gene lists of genes with pooled occupancy signal changes (Fresh = gains; Archival = losses) in the FAIRE and MNase assays from pools of three laboratory mice.

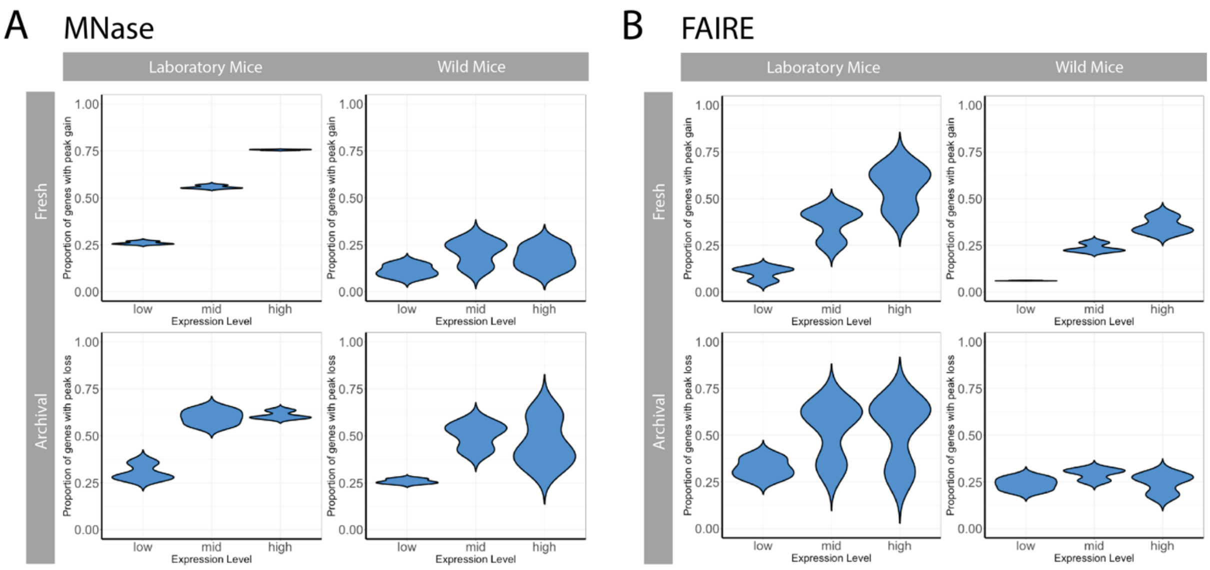


*Figure S4. Gene expression influences MNase signal detection in fresh and archival tissues*

Binning genes by expression level expressed as zFPKM (low < -2; -2 > mid < 2; high > 2), we calculated the number of genes with significant (FDR < 0.05, log10Pval < -6) peak gains (fresh tissue) or losses (archival tissues) within 2 kb upstream of the TSS. Shown are violin plots for **(A)** MNase assay signal and **(B)** FAIRE from fresh and archival liver tissue among three biological replicates in laboratory and wild-caught mice. Each individual's proportions were calculated binning genes according to their individual gene expression data.

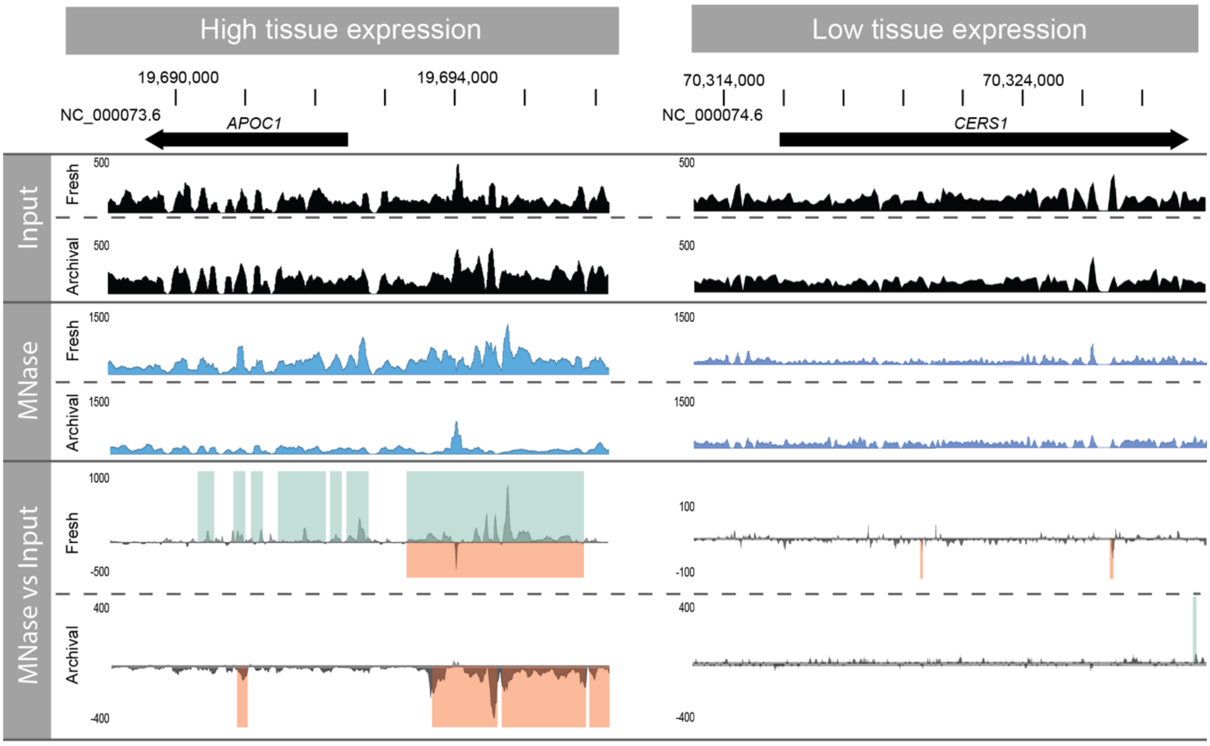


*Figure S5. Expression dependent occupancy in mouse liver*

Pooled occupancy values as wiggle traces (DANPOS3 dpeak function) for input (black) and MNase (blue) as well as differential MNase signal over input control (grey) for fresh and archival *Mus musculus* liver tissue. Occupancy values and signal changes are shown upstream of a gene highly expressed in liver (APOC1, FPKM = 38,660) and with low expression in liver (*CERS1*, FPKM = 0) as measured by RNA-Seq analysis of fresh tissue. Green and orange shading upon the differential signal panel represent significant (FDR < 0.05, log10Pval < -6) peak gains or losses across three individuals detected by DANPOS3.

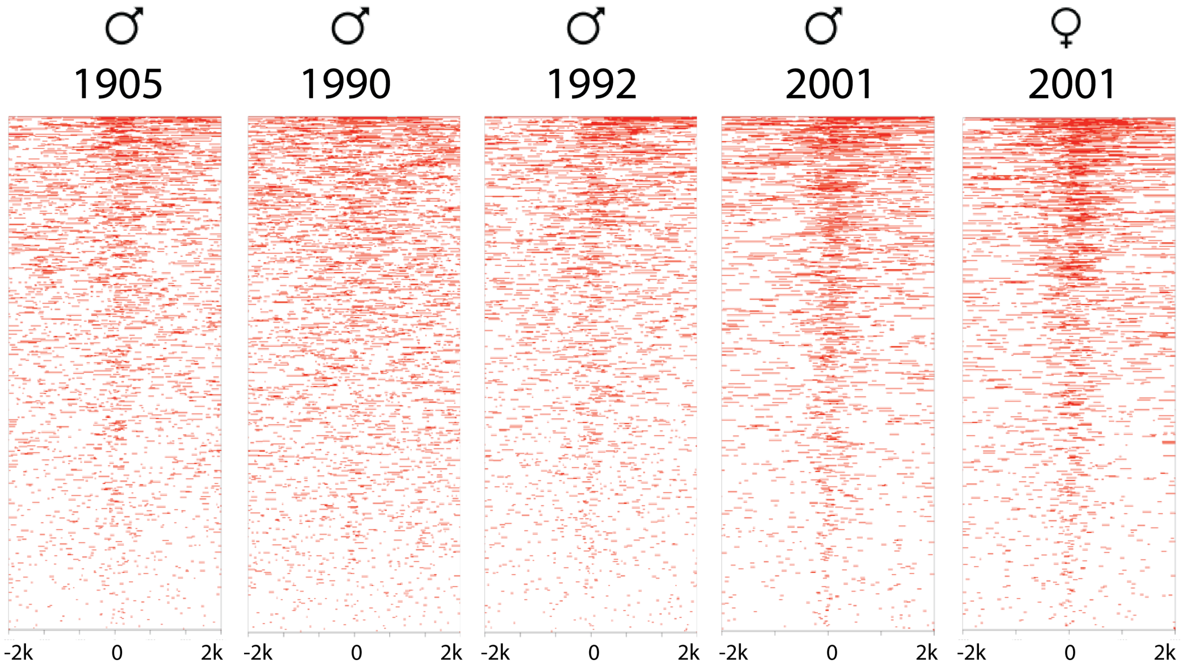


*Figure S6. Individual eastern water dragon heatmaps*

Heatmaps of MNase assay significant peak losses (FDR < 0.05, log10Pval < -6) in archival eastern water dragon liver tissues 2 kb either side of genome-wide transcription start sites.
